## Supplementary Figures for "Complement-mediated killing of bacteria by mechanical destabilization of the cell envelope"

**Supplementary Information**

**Supplementary Figures**

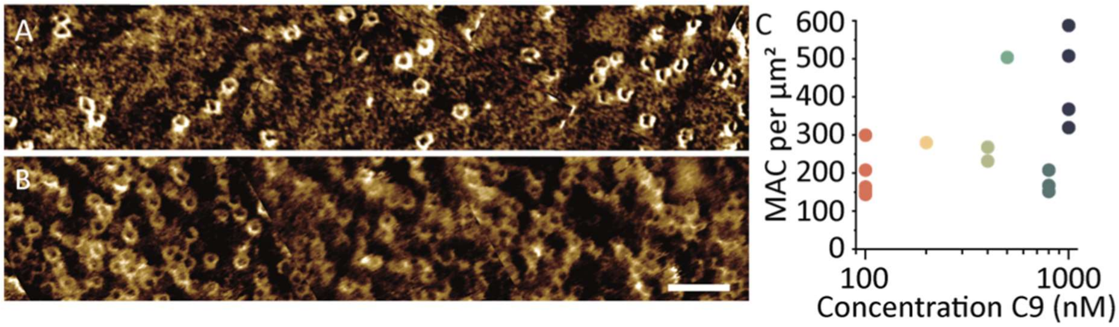


**Supplementary Fig. 1. MAC formation is highly variable between cells exposed to the same concentrations of complement proteins.**

(A-B) Whole-cell phase images show that the overall densities of MACs on the BL21 *E. coli* surface were highly varied. Some cells had (A) sparse MACs, whereas some had (B) dense packing of MACs. (C) The number of MACs in the surface was highly varied between samples, with no consistent increase for C9 concentrations ranging from 100 to 1000 nM, suggesting that the C9 concentration was not the limiting step in these experiments. Colour scale: (A) 2 deg, (B) 3.25 deg. Scale bar: 100 nm. Data refer to 16 different cells in independent experiments.

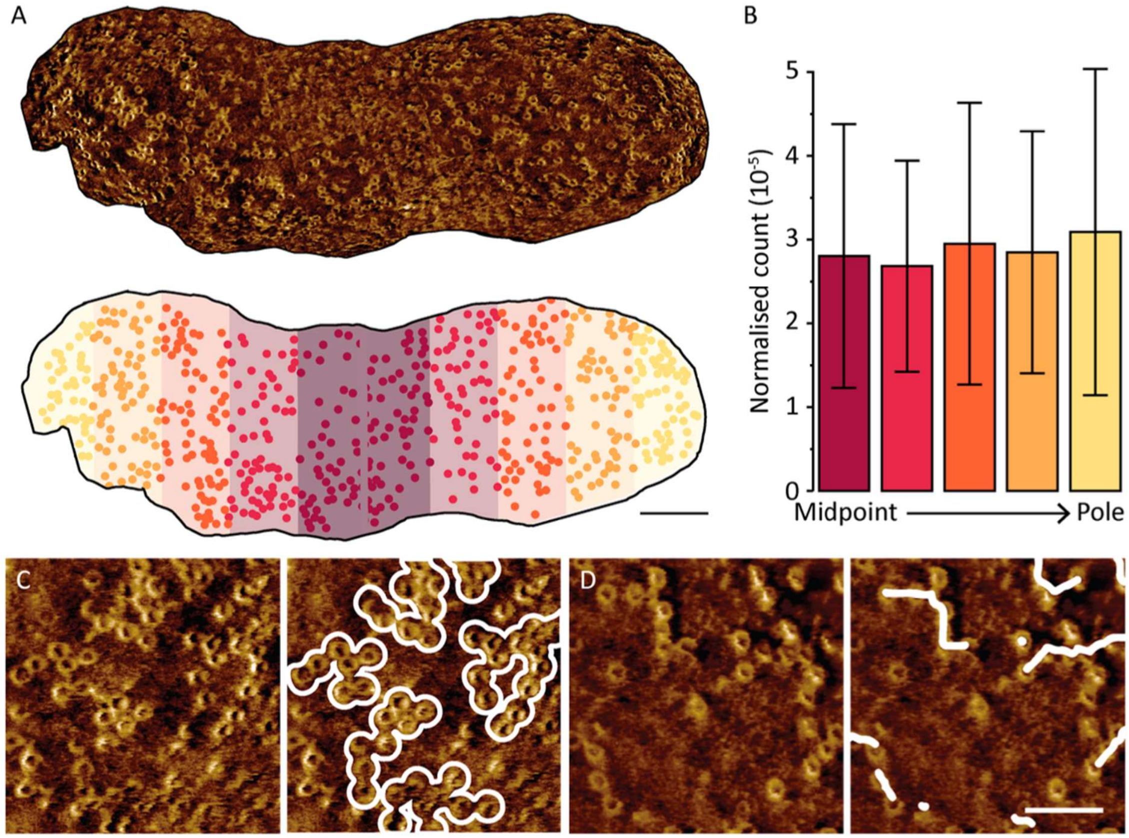


**Supplementary Fig. 2. MAC are distributed evenly over the surface of a single cell.**

(A) Top, AFM phase images were used to identify MACs and the cell midpoint. Bottom, the surface was divided into 10 regions from midpoint to pole. The numbers of MACs in each region were counted and the counts normalised to the area of each region. (B) This analysis showed the density of MACs (per $nm^2$) did not change across the cell surface for *n* = 5 different cells. Error bars indicate standard deviations (C) At first sight, MACs appeared to cluster in groups when density was high, leaving regions of bare membrane. The right image duplicates the left image, but with MAC clusters marked in white. (D) When the density was lower, MACs appeared to form in lines. The right image duplicates the left image but with branches of MACs marked in white. Colour scale: (A) 3.25 – 5 deg (C-D) 3.5 deg. Scale bars: (A) 200 nm and (C-D) 100 nm.

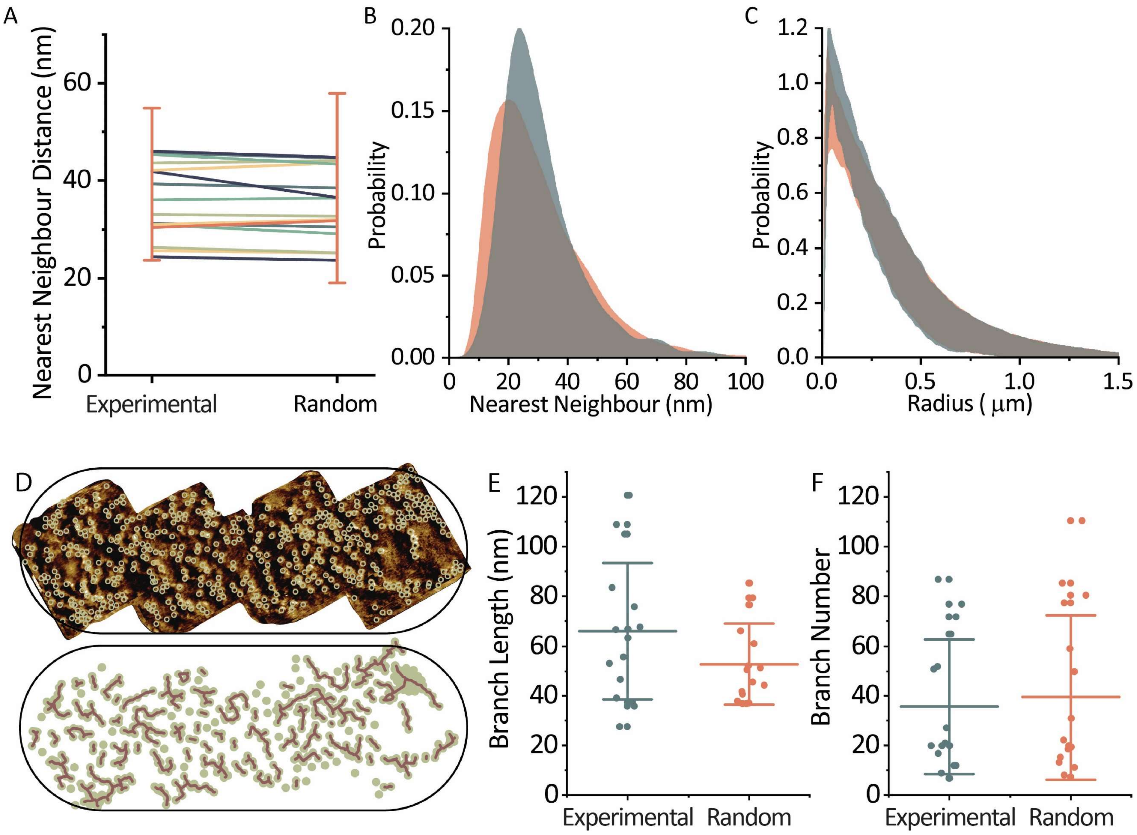


**Supplementary Fig. 3. MACs do not show significant clustering.**

From the comparison of experimental data on MAC locations with locations that were randomly generated over the same areas, the randomness of MAC distributions can be quantified. (A) Mean nearest-neighbour (MAC-MAC) distances for different samples. Error bars show typical standard deviations, only one sample standard deviation is shown for clarity. (B) Histogram of all nearest neighbours show similar distributions for real (green) and random (pink) data. (C) Radial distributions show no difference between long-distance clustering of MACs from random points. Bands corresponding to real (green) and random (pink) standard deviations are shown. (D) Chains of MACs were investigated by picking MAC points, dilating them to circles that overlap when nearby, skeletonising the resulting shape and finding the longest branch. This showed that (B) experimental and randomly generated MAC positions led to branch lengths of approximately equal size. (E) The number of branches in experimental data was also not significantly different from the random case. Data are from MACs on 16 cells.

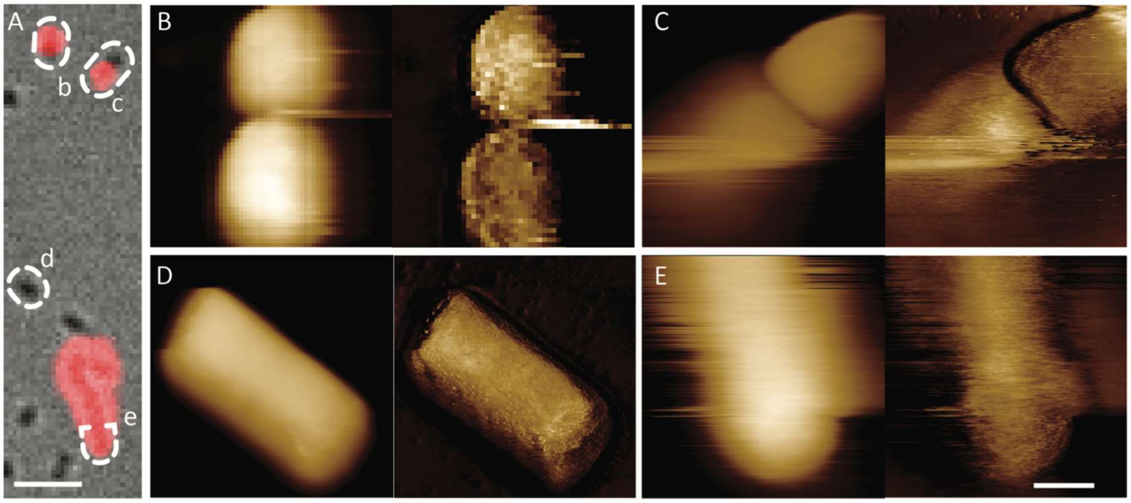


**Supplementary Fig. 4. Inner membrane permeabilization correlates with major mechanical disruption.**

(A) Merge of brightfield (grey) and SYTOX™ (red) images with bacteria imaged in B-E indicated with corresponding letters. (B-E) Height (left) and phase (right) images show that, when cells are dead (both cells in B, the bottom cell in C and the cell in E), their outer membranes become less stable. However, the membranes of live cells (top cell in C and cell in D) are stable and intact. Vertical height scale is 600 nm and phase scales are (B) 10 deg, (C) 20 deg, (D) 15 deg and (E) 15 deg. Scale bars are (A) 5 µm and (B-E) 500 nm.

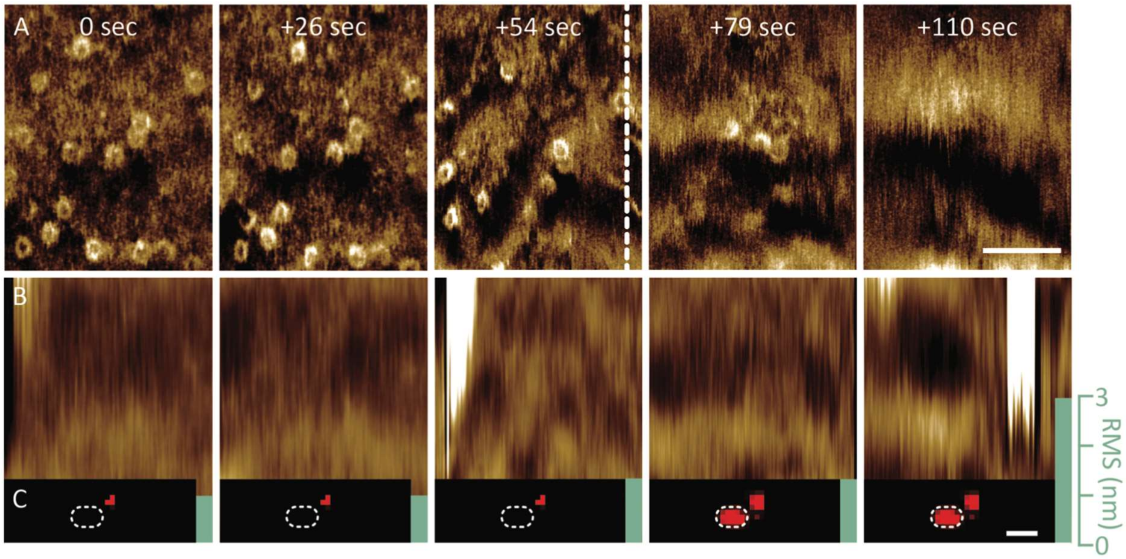


**Supplementary Fig. 5. Major disruption of the cell envelope precedes cell lysis.**

(A) Phase and (B) height images of the same area over time. (C) Fluorescence shows bacterial cell death and the green bar shows RMS roughness of height images. The start time of each scan is shown above. The AFM images are shown with the fast scan direction vertically. So, as time proceeds, subsequent scan lines are added to the image from left to right in this representation. MACs initially appear stable in phase images (0, + 26 sec), but next appear to rearrange or disappear (+54 sec), whereas in the height images, the RMS corrugation increases, after which the bacteria appear SYTOX™ positive, indicating inner membrane permeabilization. The exact time of the first SYTOX™ flush is shown by the dashed white line. Finally, the outer membrane completely destabilizes in less than 1 minute, shown by a total loss of resolution and large increase in RMS. Colour scales: (A) 2 deg and (B) 10 nm. Scale bars: (A) 100 nm and (C) 500 nm. Time points are relative to the initial image.

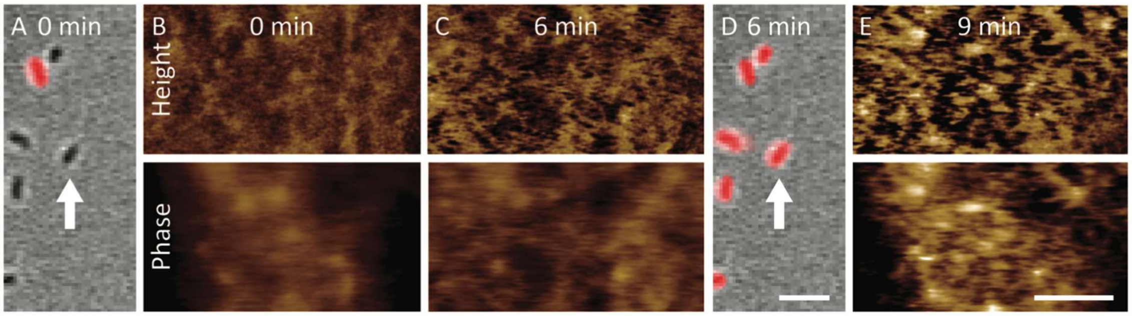


**Supplementary Fig. 6. Major outer membrane disruption or destabilization is not a generic feature of cell lysis.**

(A and D) Merges of brightfield (grey) and SYTOX™ (red) images show the imaged bacterium, indicated by the white arrow. (B, C and E) Phase and height images show that the outer membrane remains intact throughout melittin killing as the high resolution and roughening surface features are still visible. The roughness is reflected by an increase in RMS from 5 nm in B to 13 nm in C and 40 nm in E. Colour scales are 2 deg and 10 nm. Scale bars are (A and D) 5 µm and (B, C and E) 100 nm. Time points are relative to the initial image.

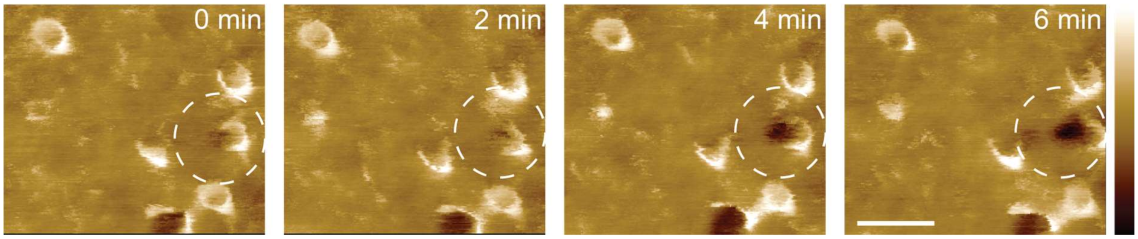


**Supplementary Fig. 7. Defect formation initiated at MAC pore.**

Sequence of AFM (height) images of BL21 *E. coli*, taken at 2 minutes per frame, showing a defect (black hole, marked by a dashed circle) initiating at a single MAC assembly.  Colour scale (bar shown on right hand side) is 20 nm. Scale bar is 50 nm**.**

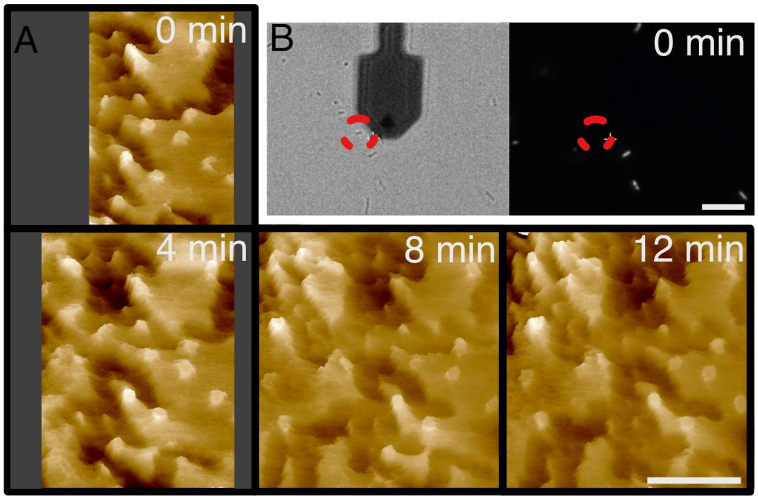


**Supplementary Fig. 8. Complement exposure leads to extensive and progressive defect formation at the bacterial surface.**

(A) Sequence of AFM (height) images of BL21 *E. coli*, cropped and aligned from Supplementary Video 2, showing MAC pores and larger (> 50 nm wide) defects in the outer membrane. Times are referenced with respect to the first recorded high-resolution image (0 min). (B) Brightfield (left) and SYTOX™ fluorescence (right) microscopy images of the bacteria in the AFM experiments, with red dashed ellipses indicating the (SYTOX™ negative) cell on which the AFM sequence was recorded. The dark paddle is the AFM cantilever. Colour scale: 30 nm; scale bars: 100 nm (A) and 10 µm (B).

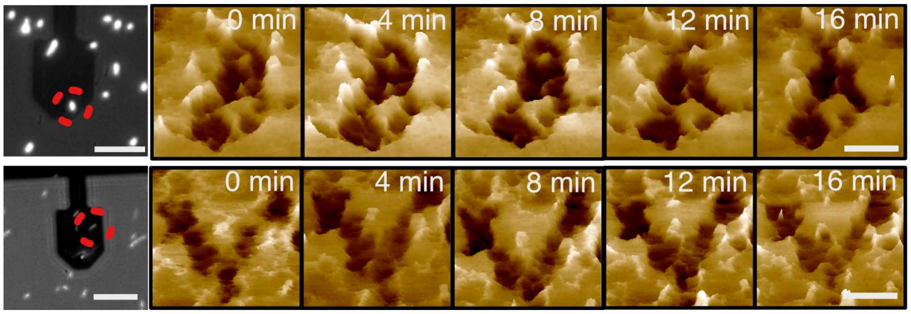


**Supplementary Fig. 9. Defects do not propagate after bacterial cell death.**

Two examples (top and bottom row) of high-resolution SYTOX™ fluorescence (first frame) and sequence of AFM images (at surface of cell marked with red dashed circle in first frame) on cells with compromised inner membranes (positive SYTOX™ signal). Colour scale: 30 nm; scale bars: 10 µm (fluorescence images); 100 nm (AFM).

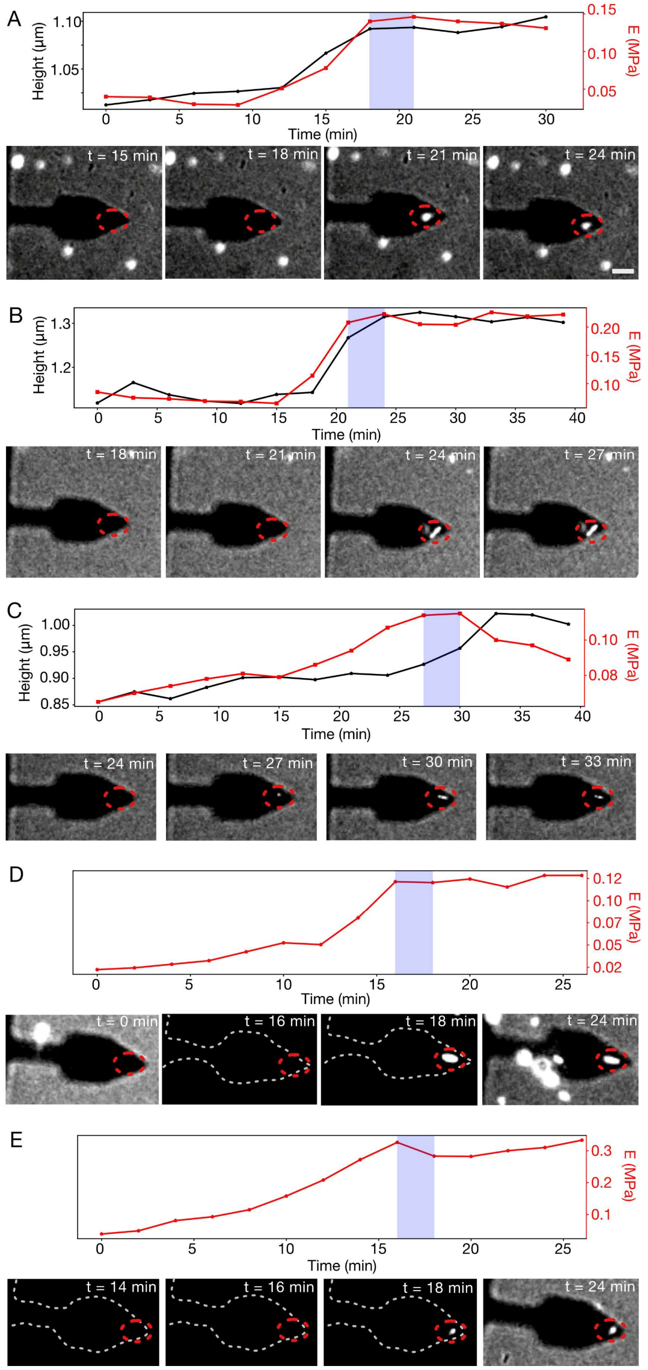


**Supplementary Fig. 10. Changes in height and surface stiffness for individual BL21 *E. coli* cells exposed to complement.**

(A-E) Changes for $n = 5$ individual cells of BL21 *E. coli*, as summarized in Fig. 4, with according SYTOX™ fluorescence data. The measured cells are marked by red, dashed circles. Scale bar: 10 µm.

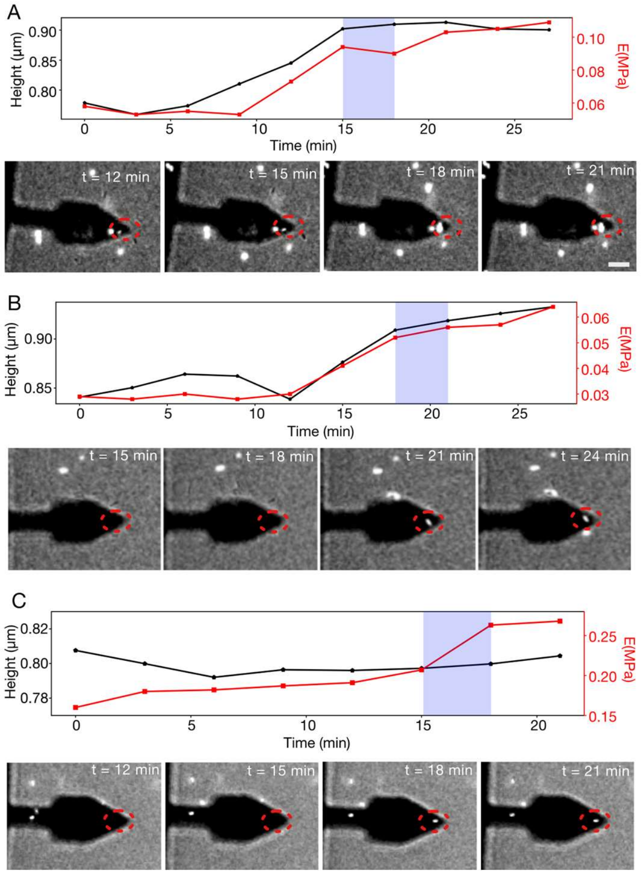


**Supplementary Fig. 11. Changes in height and surface stiffness for individual MG1655 *E. coli* cells exposed to complement.**

(A-C) Changes for *n* = 3 individual cells of MG1655 *E. coli*, as summarized in Fig. 4, with according SYTOX™ fluorescence data. The measured cells are marked by red, dashed circles. Scale bar: 10 µm.

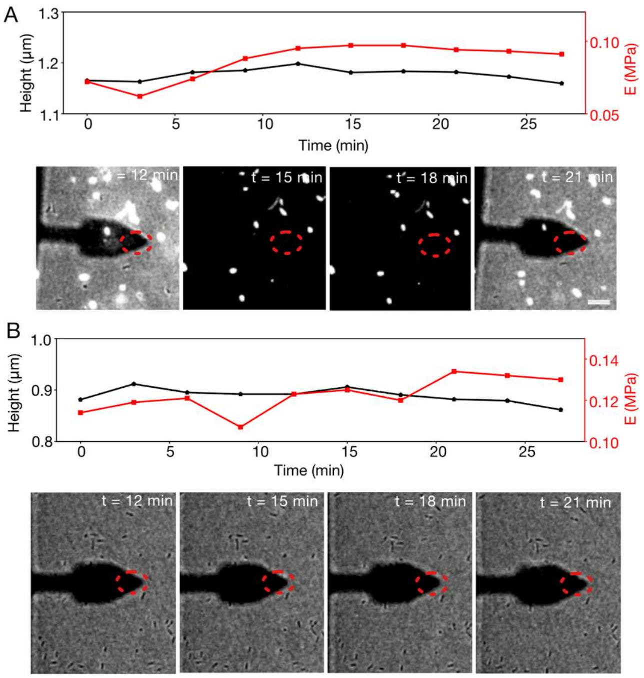


**Supplementary Fig. 12. C8 and C9 are required for complement to induce changes in height and surface stiffness of *E. coli*.**

Negative control experiments for the results shown in Fig. 4, Supplementary Figs. 10, 11). BL21 (A) and MG1655 (B) subjected to the same protocol as in (Fig. 4, Supplementary Figs. 10, 11), but without addition of C8 and C9. Over timescales investigated, no cell death or noticeable increase in cell size or surface stiffness were observed.

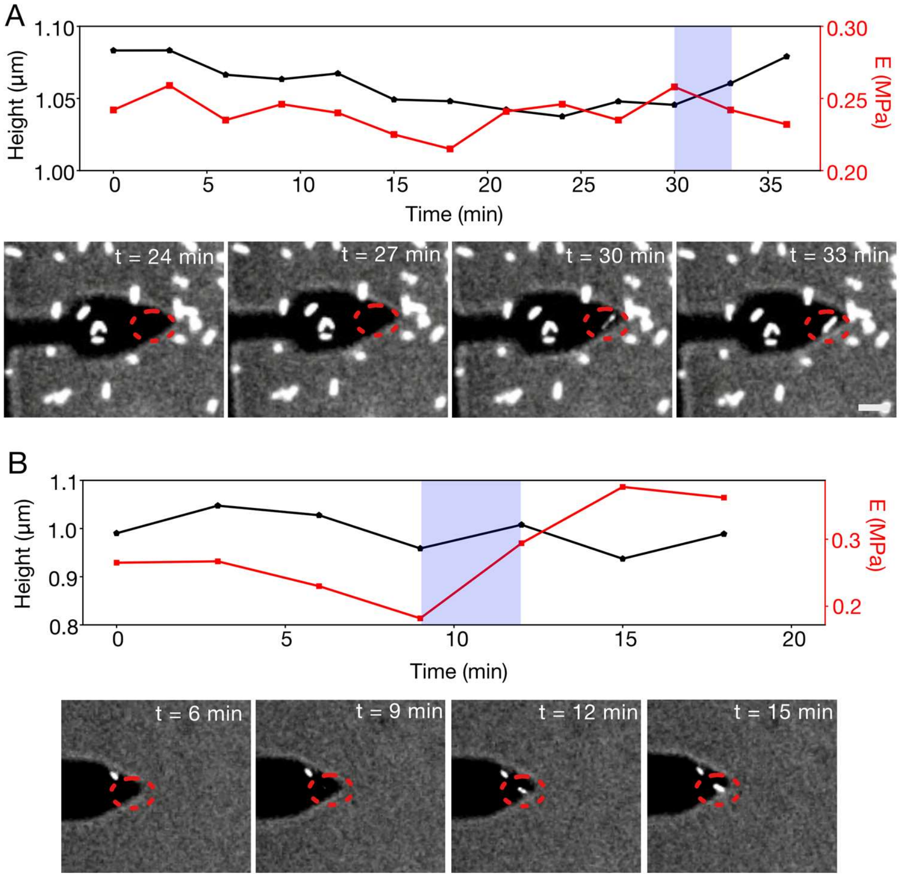


**Supplementary Fig. 13. Prior bacterial swelling and surface stiffening are not generic features of cell lysis.**

(A, B) Two negative control experiments for the results shown in Fig. 4, Supplementary Figs. 10, 11. BL21 *E. coli* is exposed to and killed by Melittin, as shown by the SYTOX$^{TM}$ staining, but show no clear trend in size or stiffness increase.

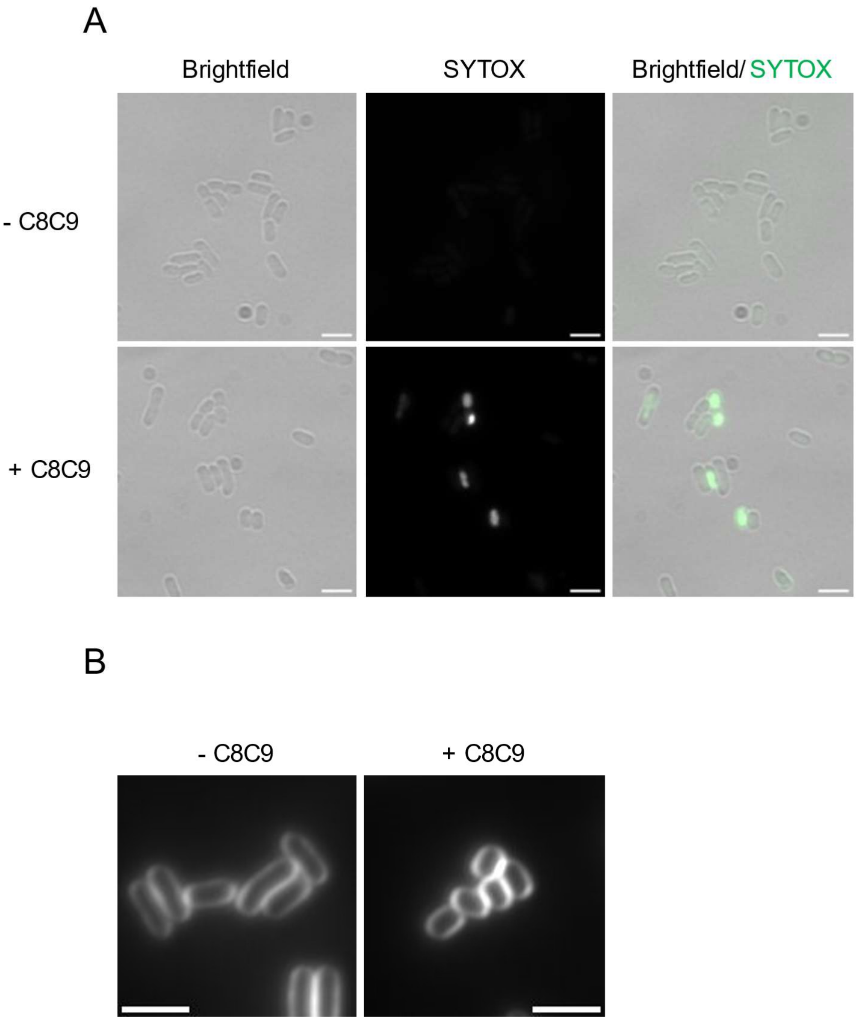


**Supplementary Fig. 14. Representative images of SYTOX™ and membrane-stained *E. coli* MG1655.** (A) Representative images of SYTOX™ stained *E. coli* MG1655 cells ± C8C9 treatment for 15 min, as used for quantitative analysis presented in Fig. 4E. (B) Representative images of FM5-95 stained *E. coli* MG1655 cells treated ±C8C9, as used for quantitative analysis presented in Fig. 4F. Scale bars: 3 µm.

### Supplementary Videos

**Supplementary Video 1.**

Complete sequence of AFM images from which crops are shown in Fig. 3B.

**Supplementary Video 2.**

Complete sequence of AFM images from which crops are shown in Supplementary Fig. 8.
